# Conserved microglial programmes, non-transferable classifiers: boundaries of the single-cell TBI molecular clock

**DOI:** 10.64898/2026.09.07.749982

**Authors:** Yan Qi, Zhang Li, Xu Xinchun, Qian Xiaobo, Li Mengge, Zhang Yingshu, Gao Rong

## Abstract

**Background:** Traumatic brain injury (TBI) pathology evolves over hours to months, and molecular staging could complement clinical and imaging-based assessment. Single-cell transcriptomics captures cell-type–resolved temporal programs, but whether a transcriptional “molecular clock” trained on one injury model can be transferred to others remains unknown.

**Methods:** We compiled four public mouse TBI single-cell/single-nucleus datasets spanning two injury models — controlled cortical impact (GSE277487: 162,892 cells; CEREBRI [GSE269748]: 39,026 nuclei) and fluid percussion injury (GSE160763: 20,642 cells) — plus an independent mild-FPI cohort (mTBI) reserved for validation (GSE247339: 85,431 cells; vehicle-treated arm of a pharmacological study), totalling 307,991 cells across four pathological stages (control, acute, subacute, chronic). Random-forest classifiers on Harmony-corrected principal components were evaluated at both cell and sample levels (held-out 10x libraries), with significance tested against a permutation null that shuffles the sample–stage mapping; we further assessed per-cell-type performance, strict leave-one-dataset-out (LODO) generalization with fully train-only preprocessing, glial-only cross-dataset transfer, and gene-level SHAP attribution.

**Results:** Within GSE277487, sample-level balanced accuracy reached 0.67 (permutation null 0.32 ± 0.09; p = 0.001), and the signal was reproduced in the independent mild-FPI cohort (0.40 vs null 0.19 ± 0.05; p = 0.0010). Cell-level cross-validation overestimated performance by ∼0.16, quantifying within-sample leakage. Cross-dataset generalization was poor (strict LODO balanced accuracy 0.20-0.31); the acute immediate-early response was untransferable (recall 0.002) and 80% of chronic (6-month) cells were assigned to the subacute class. Glial-only models raised subacute recall across models (0.91 and 0.75), but precision was indistinguishable from a subacute-only classifier (0.51 vs 0.50; 0.58 vs 0.58) and one-vs-rest AUROC was at or below chance (0.38-0.54) across four classifier families — classifier collapse, not transferable discrimination. The reproducible cross-model component was instead a gene-expression programme: a microglial complement/lysosomal damage programme (Trem2-Apoe axis) was up-regulated across the severe CCI and FPI models (13/13 concordant genes), whereas the immediate-early module was model-specific.

**Conclusions:** A single-cell molecular clock for TBI staging is feasible within datasets once within-sample leakage is controlled, but it does not transfer across datasets as a classifier: apparent cross-model transfer reflected classifier collapse onto the majority class, and the reproducible cross-model component was a microglial complement/lysosomal gene-expression programme. These findings define the boundaries of cross-dataset time inference in TBI and provide evaluation criteria for claims of transferable single-cell classifiers.

## Introduction

Traumatic brain injury (TBI) is a leading cause of death and long-term disability worldwide, with its pathological cascade evolving over hours to months after the initial insult [1]. The temporal progression — from acute excitotoxicity and neuroinflammation, through subacute glial activation and remodelling, to chronic changes that can persist for months — is central to both prognosis and treatment timing. Accurate staging of where an injured brain sits along this trajectory could therefore inform clinical decision-making, yet staging currently relies on coarse clinical and imaging surrogates rather than on the underlying molecular state.

Single-cell and single-nucleus transcriptomics have transformed our view of the TBI response by resolving cell-type–specific programmes that bulk tissue analysis cannot separate [2, 3]. Time-dependent transcriptional signatures, such as the sequential activation of microglial networks in the first week after injury [4], have been described, and transcriptomic estimates of time since injury have proven feasible in forensic contexts [5, 6]; recent syntheses have likewise outlined neuroinflammatory marker dynamics for post-traumatic interval estimation in TBI [7]. These observations raise a natural question: can a classifier trained on single-cell transcriptomes read the post-injury “molecular clock” of a cell — that is, assign it to the correct pathological stage? (Here “molecular clock” denotes transcriptomic stage inference and is distinct from epigenetic age clocks applied to brain injury [8].)

Answering this question rigorously is complicated by a methodological gap. Cells from the same 10x library are not independent: they share technical covariates, compositional effects and library-specific noise. Cross-validation at the cell level therefore partly learns library identity rather than biological time, inflating apparent accuracy. The magnitude of this within-sample leakage in single-cell classifiers is rarely quantified [9], and most TBI single-cell studies report cell-level metrics without library-aware evaluation. Similarly, whether a time-inference model trained on one injury model generalises to another — a prerequisite for any practical use — has not been systematically tested.

Here we address both questions in a systematic evaluation of four public mouse TBI datasets spanning two injury models and two assay types (single-cell and single-nucleus RNA-seq), with an independent FPI cohort (different laboratory, assay platform and brain regions) reserved for validation. We trained random-forest classifiers to assign cells to pathological stages, and evaluated them with explicit library-aware (sample-level) cross-validation, a permutation test that shuffles the sample–stage mapping, strict leave-one-dataset-out transfer with fully train-only preprocessing, and gene-level SHAP attribution. We quantify within-sample leakage directly, define which temporal programmes transfer across models and which do not, and identify the cell types and gene modules underlying the conserved component of the TBI molecular clock.

## Results

### Sample composition and cell-type architecture (Fig. 1)

We assembled four public mouse TBI single-cell and single-nucleus transcriptomic datasets spanning two injury models: GSE277487 (controlled cortical impact, CCI; 10x snRNA-seq; control, 6 h, 2 d and 4 d), GSE160763 (midline fluid percussion injury, FPI; 10x scRNA-seq; control and 7 d) and CEREBRI (GSE269748; CCI; 10x snRNA-seq; control, 24 h and 6 months, analysed as a second CCI cohort). An independent mild-FPI cohort (GSE247339; Drop-seq scRNA-seq; hippocampus and frontal cortex; sham, 24 h, 7 d and 21 d) was reserved exclusively for validation. After quality control, 307,991 cells remained for analysis (GSE277487: 162,892; GSE160763: 20,642; CEREBRI: 39,026; GSE247339: 85,431), spanning eight time points assigned to four pathological stages (control, acute 6–24 h, subacute 2–7 d, chronic 21 d–6 months). Cell-type annotation by data-driven marker scoring identified nine populations; the three primary datasets differed markedly in composition, with GSE277487 neuron-enriched (∼55% excitatory neurons) and CEREBRI and GSE160763 microglia-enriched (>50% microglia) (Fig. 1a, b).

**Figure 1.**
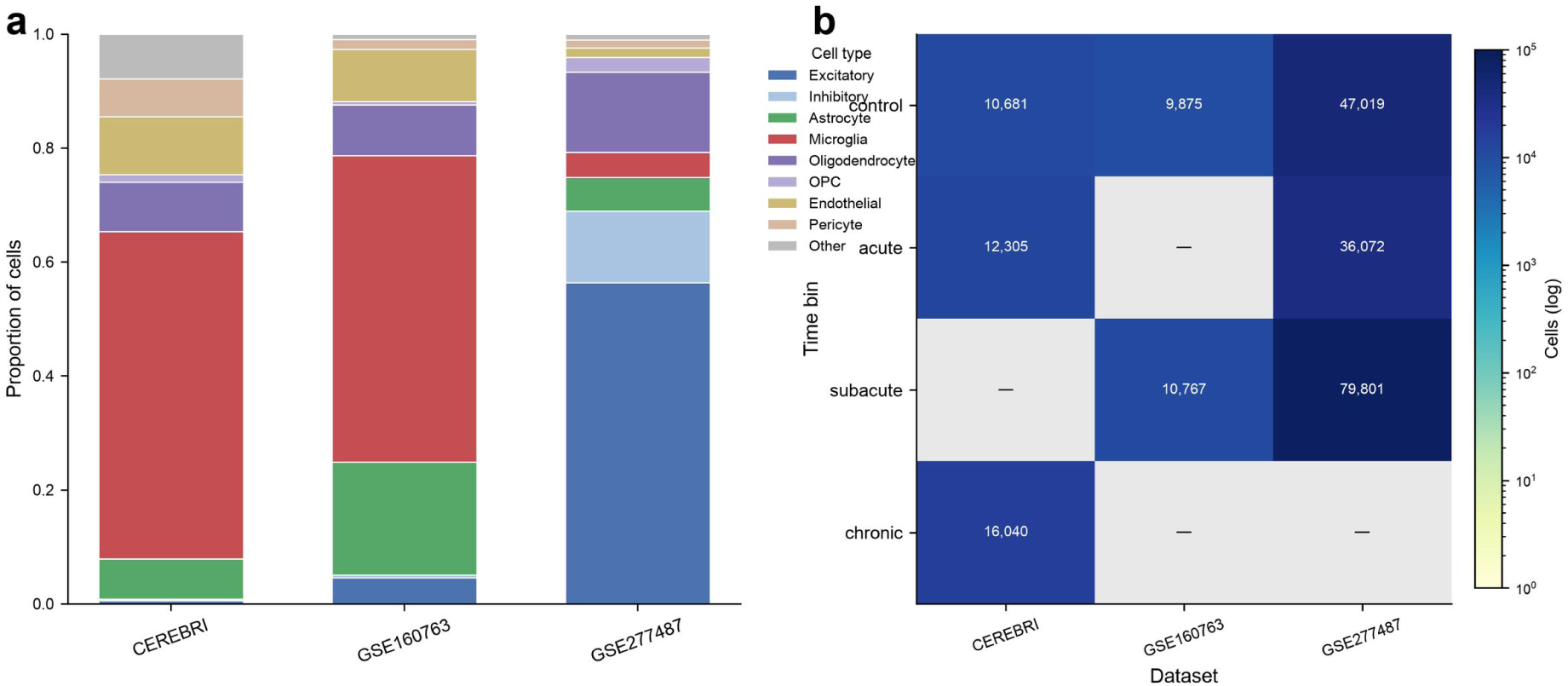
Data overview of the three mouse TBI single-cell transcriptomic datasets. (a) Proportions of the nine annotated cell types in each dataset (GSE277487, controlled cortical impact, 10x snRNA-seq; GSE160763, fluid percussion injury, 10x scRNA-seq; CEREBRI, controlled cortical impact, 10x snRNA-seq). Cell-type annotations were derived from data-driven marker scoring (Methods); GSE277487 is neuron-enriched (∼55% excitatory), whereas CEREBRI and GSE160763 are microglia-enriched (>50%). (b) Number of cells per dataset and pathological stage (control, acute, subacute, chronic), log-scaled; grey cells indicate stages absent from a given dataset. In total, 222,560 cells spanning seven time points (control; 6 h, 24 h; 2 d, 4 d, 7 d; 6 mo) were analysed. The independent mild-FPI cohort used for validation (GSE247339) is shown separately in Fig. 2c

### Sample-level within-dataset classification of post-injury stage (Fig. 2)

Because cells from the same 10x library are non-independent, we evaluated classification of post-injury stage (random forest on 30 Harmony-corrected principal components) both at the cell level and at the sample level, holding out entire 10x libraries. In GSE277487, sample-level balanced accuracy across three stages (control, acute, subacute; chance 0.33) was 0.67 (Fig. 2a). To test whether this exceeded what stage-associated technical covariates alone could produce, we permuted the sample–stage mapping across libraries 1,000 times; the observed accuracy was well above the permutation null (0.32 ± 0.09; p = 0.001; Cohen’s d = 3.74; Fig. 2b), and no permutation reached the observed value (null maximum 0.66). Cell-level cross-validation overestimated performance (0.83), indicating within-sample leakage of approximately 0.16. The combined dataset (GSE277487 + GSE160763) reached a sample-level balanced accuracy of 0.55, significant against its permutation null (0.36 ± 0.10; p = 0.043) but with a much smaller effect than the single-dataset analysis (d = 1.9 vs 3.7), a dilution we attribute to added between-dataset variance (see Discussion). In GSE277487, the subacute stage was the most accurately recovered (pooled sample-level recall 81%, versus 64% for acute and 26% for control).

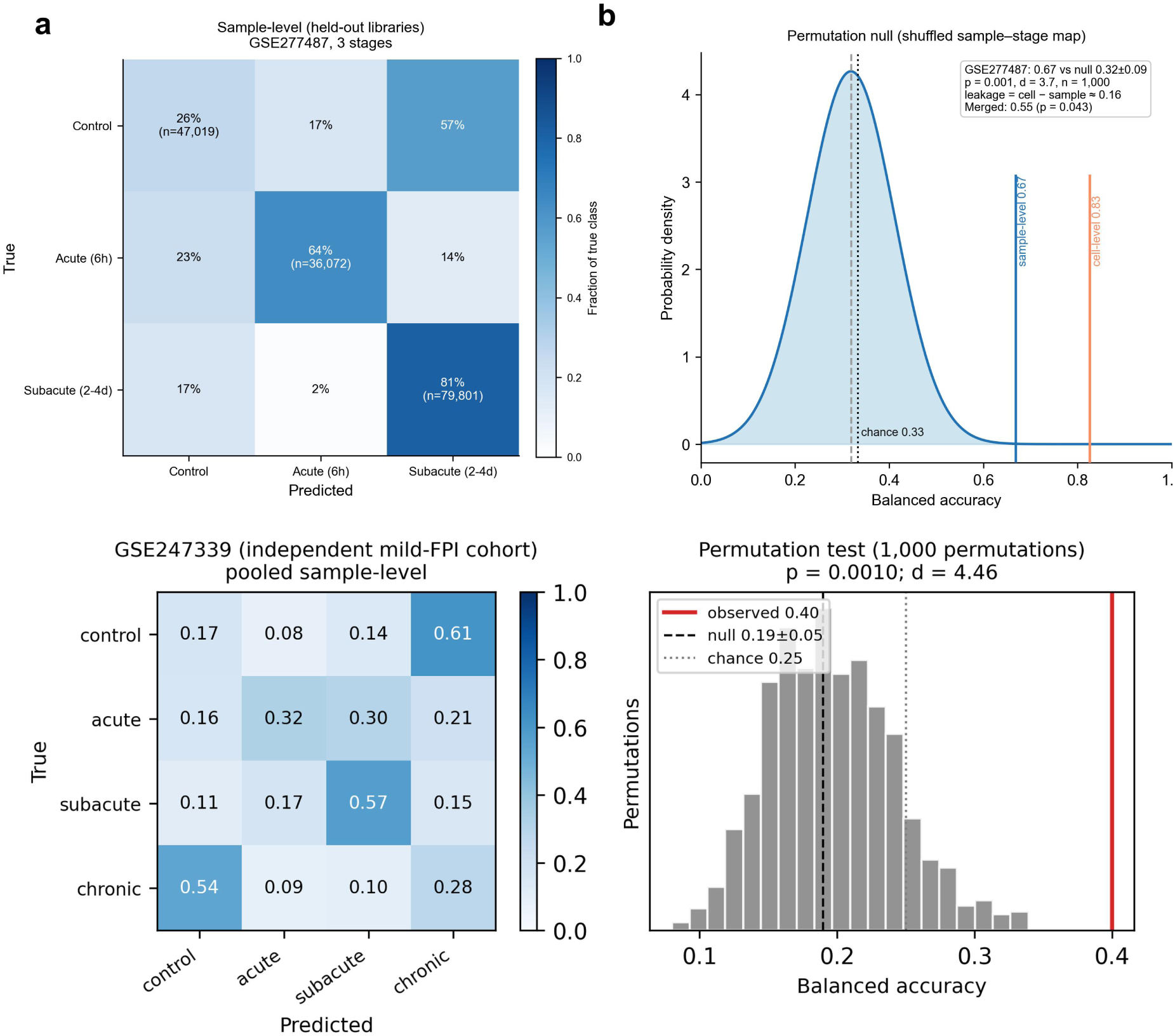
(a) Confusion matrix (row-normalized) of the random forest classifier trained on Harmony-corrected principal components and evaluated on held-out 10x libraries of GSE277487 (confusion matrix pooled across the five held-out folds; balanced accuracy was computed as the mean of per-fold balanced accuracies; three stages: control, acute, subacute; GSE277487 contains no chronic samples). (b) Three-stage GSE277487 evaluation: balanced accuracy at the cell level (0.83; within-sample leakage included) versus the sample level (0.67; held-out libraries), with the sample-level permutation null distribution shown (mean 0.32 ± 0.09; n = 1,000 permutations shuffling the sample–stage mapping); dashed lines indicate chance (0.33) and null mean. Balanced accuracy was stable across five independent split seeds (0.668–0.669; repeated sample-level CV, Methods). The gap between cell-level and sample-level accuracy (∼0.16) quantifies within-sample leakage. The combined dataset (GSE277487 + GSE160763) reached a sample-level balanced accuracy of 0.55 (permutation p = 0.043; n = 1,000). The subacute stage was the most accurately recovered (pooled sample-level recall 81%). Stages: control, acute (6 h), subacute (2–4 d). (c) Independent validation in GSE247339 (independent mild-FPI cohort; hippocampus and frontal cortex): sample-level balanced accuracy across four stages was 0.40 (confusion matrix, row-normalized, pooled across five folds), significantly above the permutation null (0.19 ± 0.05; p = 0.0010; 1,000 permutations; dashed line, chance 0.25). Stages: control (sham), acute (24 h), subacute (7 d), chronic (21 d).

### Independent validation in a second, independently profiled FPI cohort (GSE247339)

To test whether within-dataset time inference reproduces beyond the CCI cohort, we analysed GSE247339, an independent mild-FPI cohort (mTBI; vehicle-treated arm of a pharmacological study) profiled in hippocampus and frontal cortex with one library per region per animal (sham, 24 h, 7 d and 21 d; 85,431 cells after QC; 24 libraries from 12 animals), using the same marker-based annotation, feature construction and sample-level evaluation pipeline. Sample-level balanced accuracy across the four stages was 0.40 (Fig. 2c), significantly above the permutation null (0.19 ± 0.05; p = 0.0010; Cohen’s d = 4.46; 1,000 permutations), with no permutation reaching the observed value (null maximum 0.34). The absolute accuracy was lower than in the CCI cohort (0.40 vs 0.67); this may reflect the milder injury severity of this FPI cohort (a mild-TBI model), the different assay platform (Drop-seq vs 10x Chromium single-nucleus) or the different brain regions profiled, factors that cannot be fully separated. The within-dataset staging signal was nonetheless reproduced in an independent laboratory, injury cohort, assay platform (Drop-seq scRNA-seq vs 10x single-nucleus), brain region and stage scheme (four vs three stages), establishing the molecular clock as a reproducible biological phenomenon rather than a single-dataset artefact; the lower absolute accuracy further suggests that signal strength may scale with injury severity.

### The within-dataset signal is robust to model choice and feature settings

The within-dataset staging result was stable across classifiers and hyperparameters under the identical sample-level cross-validation scheme (Supplementary Fig. S1): a support vector machine with RBF kernel reached a balanced accuracy of 0.677, random forest 0.669, k-nearest neighbours 0.654 and logistic regression 0.609. Random-forest performance was insensitive to the number of trees (0.6670-0.6691 across 100-500 trees), to the number of principal components retained (0.668-0.674 across 15-30) and only modestly sensitive to the number of highly variable genes used for feature construction (0.63-0.71 across 1,000-5,000 genes under the seurat selection flavour), and repeated sample-level cross-validation with five different split seeds gave 0.6680 ± 0.0003 (GSE277487) and 0.5464 ± 0.0004 (combined dataset), so the point estimates do not depend on a particular data partition. Out-of-fold probability estimates were reasonably calibrated (Brier score 0.273 versus a three-class chance baseline of ∼0.67). In contrast, continuous regression of post-injury time from the same features failed (ridge regression R² = −0.45, RMSE = 78.4 h; random-forest regression R² = −0.20, RMSE = 43.3 h), showing that the transcriptomic signal supports stage-level but not hour-level inference.

### Temporal signal is broadly distributed across cell types at the sample level (Fig. 3)

When sample-level cross-validation was repeated within each cell type across the combined datasets (GSE277487 + GSE160763; 183,534 cells), balanced accuracy ranged narrowly from 0.49 to 0.58 across the nine populations (Fig. 3). Excitatory neurons carried the strongest signal (0.581), and no cell type showed a consistent glial advantage; the unannotated fraction fell to 0.30. These sample-level estimates contrast with cell-level estimates, in which oligodendrocytes and microglia appeared to lead — a pattern attributable to within-sample leakage rather than to cell-type–specific temporal biology.

**Figure 3.**
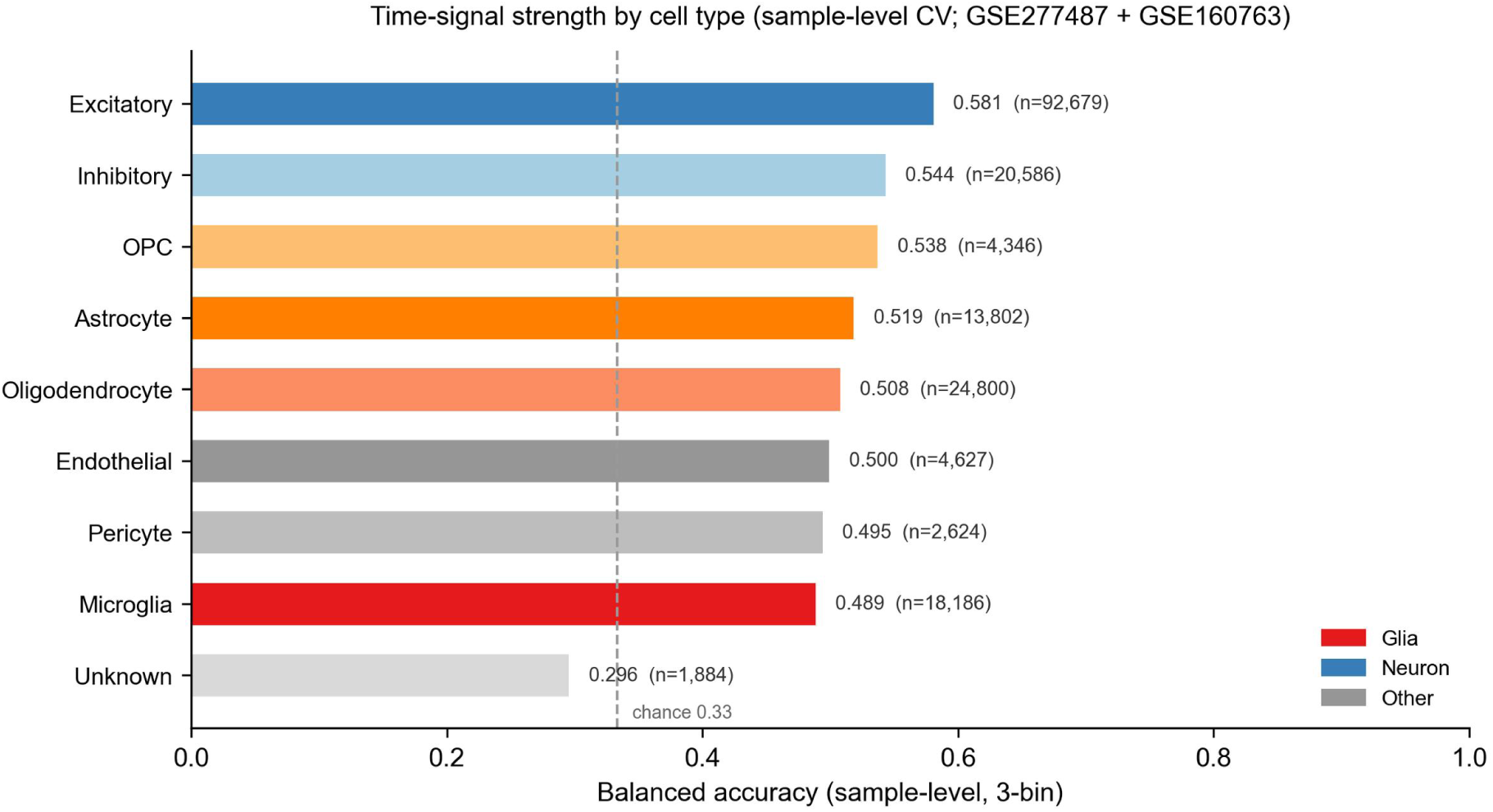
Cell-type-resolved temporal signal in the combined CCI and FPI datasets. Horizontal bars show sample-level balanced accuracy for each of the nine cell types across the combined GSE277487 + GSE160763 dataset (183,534 cells), coloured by lineage (glial: oligodendrocyte, microglia, astrocyte, OPC; neuronal: excitatory, inhibitory; other: endothelial, pericyte, unknown); per-cell-type cell counts are labelled. Dashed line indicates chance level (0.33). Temporal signal was broadly distributed across cell types (balanced accuracy 0.49–0.58), with excitatory neurons carrying the strongest signal (0.581) and no glial advantage at the sample level; unknown cells fell to 0.30.

### Cross-dataset generalization fails: classifier collapse rather than transferable discrimination (Fig. 4)

Leave-one-dataset-out evaluation with fully train-only preprocessing (HVG selection, SVD and classifier fitted on training data only) yielded balanced accuracy near or below chance for all-cell models (0.31, 0.20 and 0.21 on GSE277487, GSE160763 and CEREBRI, respectively; Fig. 4). For comparison, a “loose” scheme in which Harmony was fitted on all datasets jointly before splitting performed inconsistently — worse than the train-only scheme in two of three directions (0.06 vs 0.31 on GSE277487; 0.08 vs 0.21 on CEREBRI) but better in the third (0.31 vs 0.20 on GSE160763) — a pattern consistent with joint integration removing dataset-specific but time-informative variation. Because loose preprocessing uses the held-out data in the feature-construction step, we treat the strict train-only estimates as primary. The failure was structured: the acute immediate-early response was essentially untransferable (recall 0.002 in the GSE277487 direction; note that the acute training stage was 24 h from the snRNA-seq dataset whereas the held-out acute stage was 6 h, and immediate-early genes peak at ∼1–6 h, so part of this failure reflects a time-point scheme mismatch; direct expression analysis confirmed that the CEREBRI 24-h training libraries showed no immediate-early gene activation relative to naïve controls (mean library-level score across 15 canonical IEGs 0.90 vs 1.01; Supplementary Fig. S2a)), and chronic (6-month) cells were predominantly assigned to the subacute class (80.3%; 91.2% for glia; Supplementary Table S1). Restricting the analysis to oligodendrocytes and microglia raised subacute recall across models (0.913 and 0.748), yet balanced accuracy remained at or below chance (0.338 vs 0.333 chance on GSE277487; 0.473 vs 0.5 on GSE160763), and inspection of the confusion matrices showed that this apparent conservation was a classifier-collapse artefact: 89% and 75% of all held-out glia were assigned to the subacute class, and subacute precision (0.513 and 0.584) was indistinguishable from the precision of a classifier that assigned every cell to the subacute class (0.501 and 0.583). Consistent with this, one-vs-rest AUROC for the subacute state was at or below chance in a stricter two-dataset replication without the second CCI dataset in training — in which acute (6 h) cells were entirely unseen negative examples — (0.380 from the FPI-trained model; 0.544 from the CCI-trained model; Supplementary Table S1) — and the collapse was independent of the classifier family (random forest, logistic regression, multilayer perceptron and k-nearest neighbours all collapsed and all remained at or below chance; Fig. 4, Supplementary Table S1) and of the feature representation: re-running the same transfer on a deep generative latent space (scVI, 30 latent dimensions fitted on training data only) reproduced the collapse (97-98% of held-out glia assigned to subacute; one-vs-rest AUROC 0.45-0.54; Supplementary Table S1). A glial classifier trained on CCI likewise failed to discriminate subacute (7 d) cells in the independent mild-FPI cohort (recall 0.60 but one-vs-rest AUROC 0.367). The reverse direction was confounded by the stage-scheme mismatch: the independent FPI cohort includes a chronic (21 d) stage with no counterpart in the CCI data and assigned most CCI cells to that stage, illustrating that differing stage schemes are themselves a barrier to transfer.

**Figure 4.**
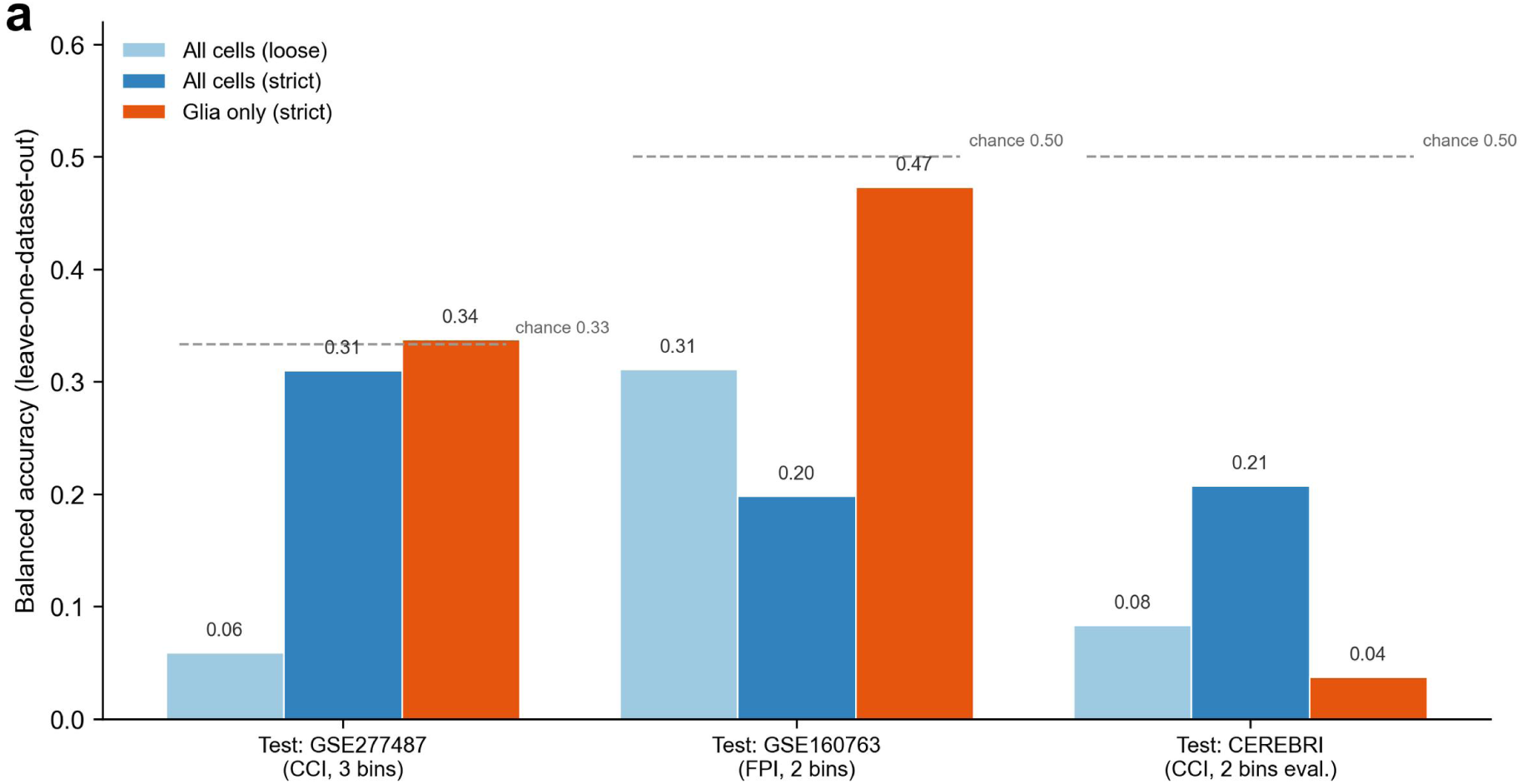
Leave-one-dataset-out (LODO) cross-dataset generalization. Balanced accuracy on each held-out dataset for three analysis schemes: all cells with loose preprocessing (Harmony fitted on all data; light blue), all cells with strict train-only preprocessing (HVG selection, SVD and random forest fitted on training data only; dark blue), and glia-only (oligodendrocytes + microglia) with strict preprocessing (orange). Held-out datasets: GSE277487 (CCI; three-stage evaluation — control, acute, subacute — chance 0.33), GSE160763 (FPI; two-stage evaluation, chance 0.50), CEREBRI (CCI snRNA-seq; two-stage evaluation of control and acute, chance 0.50; its chronic-stage cells were excluded from the metric and are reported separately in Supplementary Table S1). Glia-only models improved balanced accuracy over all-cell models in the GSE160763 direction (0.47 vs 0.20) but remained at or below chance in every direction (0.47 vs 0.50; 0.34 vs 0.33; CEREBRI ≤ 0.04), and all models failed on CEREBRI (≤0.21 for all-cell). The high subacute recall of the glia-only models (0.91 and 0.75) reflects classifier collapse onto the subacute class rather than transferable discrimination: subacute precision equalled the subacute-only baseline, and one-vs-rest AUROC was at or below chance in both directions and across four classifier families (Supplementary Table S1).

### Gene-level attribution implicates neuroinflammatory and injury-response modules (Fig. 5)

Gene-space SHAP attribution of the within-dataset classifier highlighted five coherent modules: complement/innate immunity (C1qa/b/c, Tyrobp, Hexb), myelin/glial structure (Mbp, Plp1, Gfap), lipid/cholesterol metabolism (Apoe, Clu, Ptgds), iron metabolism/oxidative stress (Ftl1, Mt1/Mt2) and immediate-early genes (Egr1, Fos) (Fig. 5); highly abundant mitochondrial and structural transcripts (e.g. Lars2, Malat1) also ranked at the top by absolute SHAP value but lack stage-specific patterning. Stage-resolved expression in GSE277487 specified the direction of these attributions: immediate-early genes (Egr1, Fos, Fosb, Junb) were most highly expressed at the acute (6 h) stage, whereas complement/innate-immunity (C1qa/b/c, Tyrobp, Hexb), lipid/cholesterol (Apoe, Clu, Ptgds), iron-metabolism (Ftl1) and astrocytic (Gfap) genes were progressively elevated at the subacute (2–4 d) stages (Supplementary Fig. S2), consistent with the time-dependent engagement of the classical complement pathway after TBI reported at the protein level [18]. These patterns map onto the principal pathological cascades of TBI and support a biological, rather than purely technical, basis for the temporal signal; they also resolve the apparent tension between the high importance of immediate-early genes and their poor cross-dataset transfer, which reflects experiment-specific induction of this module. At the gene-expression level, the reproducible cross-model component of the subacute response was the complement/innate-immunity and lysosomal microglial programme: a damage-associated microglia (DAM)-like core (Trem2, Apoe, Ctsb/d/l/z, Lyz2, Spp1, Cd9, Gpnmb, Itgax, Lpl, Timp2) was up-regulated in microglia of both the CCI (2–4 d) and the FPI (7 d) datasets relative to their controls, with 13/13 genes concordant across the two models (library-level mean expression; expression-matched permutation null, p = 0.004; Supplementary Table S3), and the full programme separated subacute from control libraries of the CCI cohort with a library-level effect size of d = 3.2. This programme was not detected in the mild-FPI cohort at any sampled time point under that cohort’s design (sham animals harvested at the 21-day time point only; Supplementary Table S3). The reproducible cross-model element of the TBI subacute response is therefore a microglial gene-expression programme rather than a transferable classifier state.

**Figure 5.**
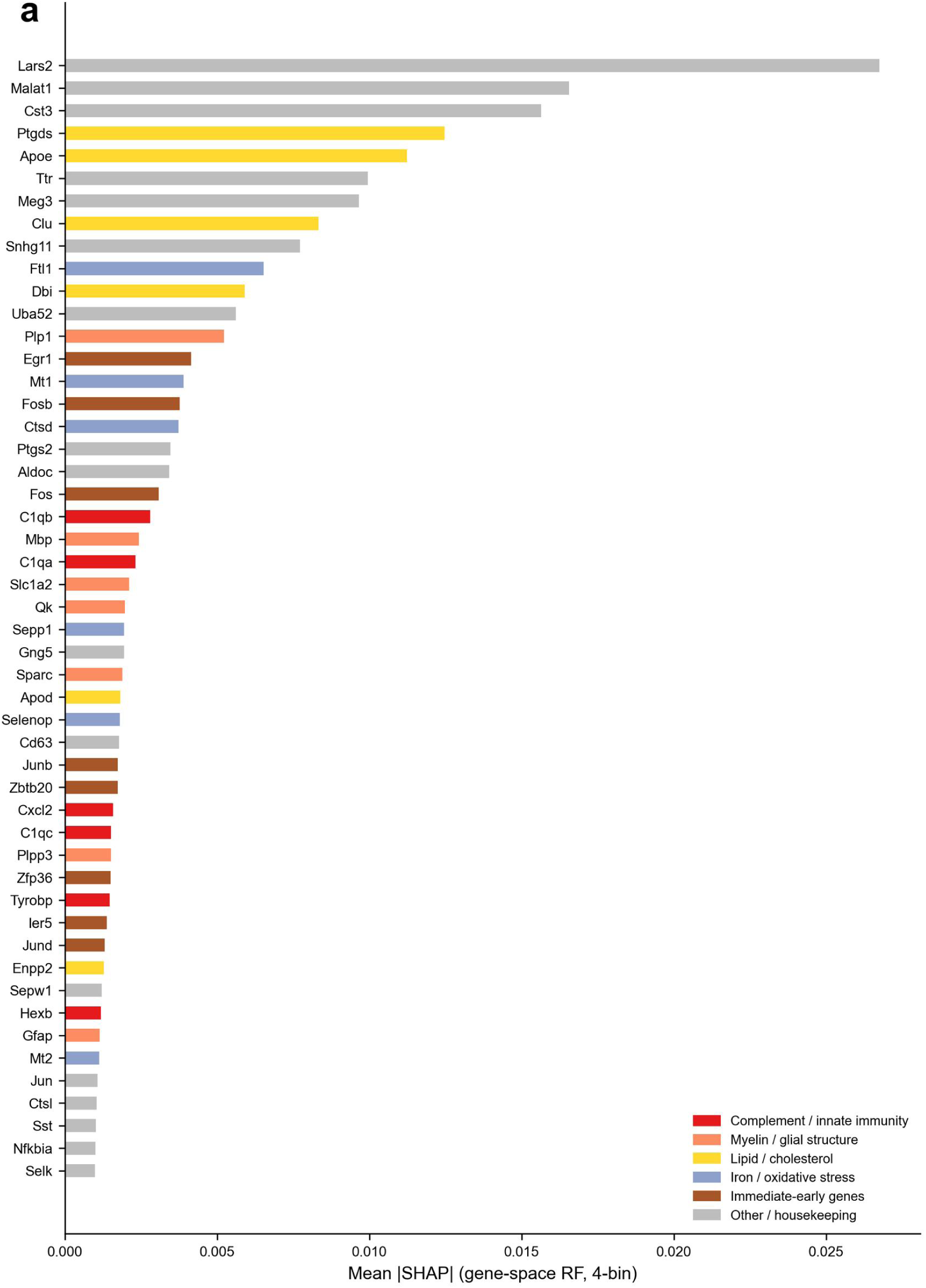
Gene-level SHAP attribution for the four-time-bin classifier within GSE277487 (control, 6 h, 2 d and 4 d). Horizontal bars show mean |SHAP| values from the gene-space random forest (maximum depth 15, 1,500-cell subsample) for the top 50 genes. Colours denote functional modules: complement/innate immunity (C1qa/b/c, Tyrobp, Hexb; red), myelin/glial structure (Mbp, Plp1, Gfap, Slc1a2; light orange), lipid/cholesterol (Apoe, Clu, Ptgds, Dbi; yellow), iron/oxidative stress (Ftl1, Mt1/Mt2, Ctsd; light blue), immediate-early genes (Egr1, Fos/Fosb, Junb; brown) and other/housekeeping genes (grey).

## Discussion

We show that post-injury stage can be inferred from single-cell transcriptomes of mouse TBI tissue, and that the reliability of such inference is substantially lower than cell-level cross-validation suggests once the non-independence of cells within 10x libraries is taken into account. In GSE277487, sample-level classification of three pathological stages reached a balanced accuracy of 0.67, significantly above a permutation null that shuffles the sample– stage mapping (0.32 ± 0.09; p = 0.001), while cell-level evaluation inflated performance by ∼0.16. This leakage is not an artefact of our pipeline: it arises because cells from the same library share technical and compositional covariates, so that naive cross-validation partly learns library identity rather than biological time. The ∼0.16 gap we quantify implies that cell-level metrics in single-cell classifiers should be read as upper bounds and that library-aware evaluation should become standard practice in this growing literature [9].

The markedly weaker sample-level signal in the combined dataset (0.55; p = 0.043; d = 1.9), relative to GSE277487 alone (0.67; d = 3.7), calls for an explanation. Two factors contribute. First, GSE160763 contributes only two stages (control and 7 d) and few libraries, so the permutation null for the combined analysis is both higher (0.36 vs 0.32) and more dispersed than for GSE277487 alone. Second, merging a second model with a different injury device, severity and tissue preparation adds between-dataset variance that a three-stage grouping cannot absorb. We therefore regard the GSE277487 analysis as the primary within-dataset evidence and the combined analysis as a conservative robustness check; the effect remained significant (p = 0.043) despite the dilution. This pattern also cautions against pooling heterogeneous single-cell datasets for time-inference tasks without explicit library-level controls.

At the sample level, temporal signal was not dominated by any cell type (balanced accuracy 0.49–0.58 across nine populations), in contrast to cell-level estimates that appeared to favour oligodendrocytes and microglia — a discrepancy attributable to within-sample leakage. Cross-dataset transfer initially presented a different picture: glia-only models returned high subacute recall (0.91 and 0.75), which we first read as evidence for a conserved glial subacute programme. Systematic inspection showed this interpretation to be an artefact of classifier collapse: the models assigned 89% and 75% of all held-out glia to the subacute class, subacute precision was indistinguishable from the precision of a subacute-only classifier, and one-vs-rest AUROC remained at or below chance in both transfer directions and across four classifier families (random forest, logistic regression, multilayer perceptron and k-nearest neighbours). The high recall therefore reflected the majority-class structure of the training data combined with cross-domain shift, not a transferable discrimination of the subacute state. This is a second evaluation pitfall in single-cell cross-dataset studies, complementary to the within-sample leakage quantified above: under class imbalance and domain shift, per-class recall and balanced accuracy can appear informative while carrying no transferable signal, and claims of cross-dataset generalizability should be supported by precision relative to a majority-class baseline and by one-vs-rest AUROC rather than by isolated recall values. The conserved element of the subacute response did survive at the gene-expression level, where a microglial DAM-like programme (Trem2–Apoe–lysosomal; 13/13 concordant genes across the CCI and FPI models) was reproducibly up-regulated — consistent with recent reports of TREM2-dependent DAM-like microglial states after TBI [14, 15] and of TREM2-dependent microglial phagocytosis in the subacute phase [15] — whereas the neuronal immediate-early response was not conserved, consistent with its time-point-specific (1–6 h) induction and the mismatch between the 24-h acute training stage and the 6-h held-out acute stage. The dissociation between dataset-specific classifier states and a reproducible gene-expression programme is itself informative: it suggests that what transfers across TBI models is not a decision boundary over cells, but the underlying molecular programme, and that claims about conserved cell states should be anchored in the latter.

A broader context for the negative transfer result is that cross-dataset transferability depends on the existence of a shared reference. Integrative atlases succeed when datasets can be anchored to a common reference state space: for example, 36 neural-organoid datasets spanning 26 differentiation protocols were integrated by projecting onto a developing-human-brain reference with scArches, yielding a reusable scaffold for annotating new protocols and disease models [19]. The TBI datasets examined here lack such an anchor: each injury model defines a distinct pathophysiological state space (severity, time window, injury physics), and no time-resolved TBI reference atlas exists onto which new datasets could be projected before classification. Our observation that transfer fails across classifier families and across representations, including a deep generative (scVI) latent space, is therefore consistent with the view that cross-dataset transfer is not a property of the model but of the availability of a shared, well-annotated reference; building such a reference for TBI time inference is a prerequisite for transferable staging, and the evaluation metrics introduced here (precision relative to a majority-class baseline, one-vs-rest AUROC) are the criteria by which such a reference should itself be validated.

The within-dataset staging signal was not confined to the CCI cohort: in GSE247339, an independent mild-FPI cohort (vehicle-treated arm of a pharmacological study [13]) profiled by a different laboratory in hippocampus and frontal cortex, sample-level balanced accuracy across four stages was 0.40, significantly above its permutation null (0.19 ± 0.05; p = 0.0010; 1,000 permutations). The lower absolute accuracy (0.40 vs 0.67) may reflect the milder injury severity of this cohort, the different assay platform (Drop-seq) or region composition; although these factors cannot be fully separated, the reproducibility of a significant within-dataset signal across laboratories, assay platforms and brain regions supports the biological basis of the staging signal rather than a single-dataset artefact. The validation design is therefore layered: cross-model transfer is tested between CCI and an independent midline-FPI dataset (GSE160763), whereas GSE247339 provides within-model (FPI) replication at a milder severity from a different laboratory, platform and brain regions. The absence of a detectable DAM-like programme in this cohort is not evidence against microglial TREM2 responses in mild injury per se — a TREM2-positive microglial phenotype has been described after mild TBI [17] — and instead delimits the sensitivity of the present cohort design (single 21-d sham time point, Drop-seq platform, cortex and hippocampus pooled).

The assignment of 80% of chronic (6-month) cells to the subacute class must be interpreted with particular care, because the LODO classifiers showed a general preference for the subacute class (89% and 75% of all held-out glia, including genuinely control and acute cells, were assigned to subacute). Part of the chronic-to-subacute assignment is therefore a classifier-collapse artefact rather than a biological signal. After this is taken into account, the residual possibility of a genuinely persistent subacute-like state remains open, and is supported by reports of sustained neuroinflammation and gliosis months after experimental TBI [10, 12]; alternatively, batch or protocol differences between the chronic dataset (CEREBRI; snRNA-seq, distinct laboratory and tissue-preparation pipeline) and the training datasets may contribute. Our design cannot separate these contributions, and we state this limitation explicitly. If a persistent component exists, the finding would argue that “chronic” TBI tissue retains elements of an active inflammatory programme rather than a purely quiescent state — a hypothesis testable in dedicated longitudinal experiments.

Several limitations merit emphasis. First, although the within-dataset staging signal was independently reproduced in a second FPI cohort (GSE247339; 0.40, p = 0.0010), the analyses still rest on a limited number of libraries (12 in GSE277487; 24 in GSE247339), confidence intervals around per-fold accuracy are wide, and the lower absolute accuracy in this cohort underscores that staging across laboratories, platforms and severities will require larger cohorts. In GSE247339, cortex and hippocampus libraries were obtained from the same animals, so library-level cross-validation does not fully exclude within-animal correlation between regions; however, grouping the two region libraries of each animal (animal-level cross-validation over 12 animals) retained a significant staging signal (balanced accuracy 0.35 vs permutation null 0.126 ± 0.049, p = 0.0099, 100 permutations; Supplementary Table S4), indicating that the signal is not attributable to within-animal correlation between regions. Sham animals were harvested at the 21-day time point, so the control stage reflects a single post-surgery time. Second, stage labels are derived from sampling time, not from pathophysiological verification, so misclassification of true biological stage is possible. Third, CEREBRI cell-type annotations proved unreliable against marker-based evaluation, and we therefore used data-driven annotation throughout; this discrepancy itself illustrates the need for annotation validation in multi-dataset analyses. Fourth, all data are from mouse models; transfer to human TBI will require human single-cell atlases with time-stamped sampling. Fifth, our permutation test, while correctly destroying the sample–stage mapping, preserves within-library structure and therefore estimates the null for the library-grouped design specifically.

Our results define both the feasibility and the boundaries of single-cell time inference in TBI: a molecular clock is detectable within a dataset once leakage is controlled, but it does not transfer across datasets as a classifier — apparent cross-model transfer reflected classifier collapse onto the majority class (a pitfall we show to be detectable by precision and one-vs-rest AUROC), and the reproducible cross-model component was a microglial DAM-like gene-expression programme rather than a transferable classifier state. We suggest that library-aware evaluation, permutation testing, annotation validation and majority-class-aware metrics should become standard practice in single-cell time-inference studies, and that claims of cross-dataset generalizability be accompanied by explicit boundary statements and, where possible, by anchor in gene-expression programmes rather than in classifier states.

## Methods

### Datasets and quality control

We analysed four public mouse TBI datasets: GSE277487 (controlled cortical impact, CCI; 10x Chromium snRNA-seq; control, 6 h, 2 d, 4 d; 162,892 cells after QC), GSE160763 (midline fluid percussion injury, FPI; 10x scRNA-seq; control and 7 d; vehicle-treated samples of a microglia-depletion study; 20,642 cells) [10], CEREBRI (GSE269748; CCI; 10x snRNA-seq; control, 24 h, 6 mo; 39,026 nuclei) [2] and GSE247339 (mild fluid percussion injury, mild FPI; Drop-seq scRNA-seq; hippocampus and frontal cortex, one library per region per animal; vehicle-treated sham and TBI arms of a pharmacological (thyroxine) study [13]; sham, 24 h, 7 d, 21 d; 85,431 cells after QC). Raw count matrices were downloaded from GEO (GSE277487, GSE160763, GSE247339 and GSE269748 [CEREBRI]). PLX5622-treated samples in GSE160763 and the damaged sample N_1 were excluded, retaining the vehicle-treated sham and TBI (7 d) samples. GSE247339 was used exclusively as an independent validation dataset and was not included in any training or feature-construction step of the primary analyses; sham samples served as the control stage, and 24 h, 7 d and 21 d samples were assigned to acute, subacute and chronic stages, respectively. For all datasets, cells with fewer than 200 detected genes or more than 20% mitochondrial reads were removed. Counts were normalised to 10,000 per cell and log1p-transformed.

### Pathological stage definition

Based on the reported sampling times, cells were assigned to four pathological stages: control (0 h), acute (6–24 h), subacute (2–7 d) and chronic (6 months). GSE277487 contains control, acute and subacute stages only (no chronic samples); CEREBRI contributes the chronic stage.

### Cell-type annotation

Cell types were annotated by data-driven marker scoring rather than using the official annotations, because the official CEREBRI “Glutamatergic_neuron” subset showed a microglia-dominated expression profile (C1QB detected in ∼50% of cells vs. SLC17A7 in ∼3%), indicating that official labels are not reliable as ground truth in this dataset; CEREBRI is the only dataset of the four whose deposition includes an official cell-type annotation, the remaining datasets being deposited as raw count matrices, and the same data-driven procedure was therefore applied uniformly. For each of nine cell types, the mean normalised expression of curated positive markers was scored per cell; cells were assigned the type with the highest mean positive-marker score; cells in which every marker score was zero were labelled Unknown. Marker sets comprised canonical genes for excitatory/inhibitory neurons, oligodendrocytes, microglia, astrocytes, OPCs, endothelial cells, pericytes and unknown cells (full lists in Supplementary Table S2).

### Feature representation

The 3,000 most highly variable genes (Seurat v3 flavour) were scaled to unit variance (maximum 10) and projected onto 50 principal components. To remove dataset-specific technical effects, Harmony [11] was applied to the PCA embedding with dataset identity as the batch variable (20 iterations, converged after 4). The first 30 Harmony-corrected components were used as features for all classifiers. Features were always derived from the training data only in cross-dataset experiments (see below).

### Classification and performance evaluation

For within-dataset evaluation, random forest classifiers (300 trees, all other parameters default) were trained to predict the pathological stage from the 30 Harmony components. Because cells from the same 10x library are not independent, we evaluated performance at two levels: (i) cell-level cross-validation (stratified 5-fold, ignoring library structure) and (ii) sample-level cross-validation (GroupKFold with 5 groups defined by 10x library, so that all cells of a held-out library are excluded from training). Balanced accuracy (mean per-class recall over the classes present in the training set) was the primary metric; chance level is 1/3 for the three-stage evaluations (GSE277487 and the combined GSE277487+GSE160763 dataset) and 1/4 for the four-stage GSE247339 evaluation.

### Permutation test

To test whether sample-level accuracy exceeded what could be achieved by stage-associated technical or compositional covariates alone, we generated a null distribution by permuting the sample–stage mapping across libraries: each 10x library retains its stage label distribution collapsed to its modal stage, and the mapping between libraries and stages is shuffled 1,000 times (GSE277487, the combined analysis and GSE247339) while preserving the number of libraries per stage. With only 30 permutations the minimum achievable p is 0.032; we therefore used 1,000 permutations for all sample-level analyses (minimum achievable p = 0.001). For each permutation, sample-level cross-validation was repeated with random forests of 100 trees. The empirical p value was computed as (number of permutations with accuracy ≥ observed + 1) / (number of permutations + 1); Cohen’s d was computed as the difference between observed and null means divided by the null standard deviation.

For model comparison, logistic regression (L2 penalty), support-vector machines (RBF kernel) and k-nearest neighbours (k = 15) were evaluated under the same sample-level cross-validation scheme as the random forest. Hyperparameter sensitivity was assessed by varying the number of trees (100-500), the number of principal components retained (15-30) and the number of highly variable genes used for feature construction (1,000-5,000 under the seurat selection flavour; all other pipeline steps unchanged). Repeated sample-level cross-validation (five independent runs with split seeds 42-46) quantified the stability of the accuracy estimates. Probability calibration was evaluated with the Brier score and a reliability curve computed from out-of-fold predictions of the reference random forest. For continuous time inference, ridge and random-forest regressors were trained on the same features with log1p-transformed post-injury hours as the target, under the same sample-level cross-validation scheme.

### Cross-dataset (leave-one-dataset-out) evaluation

For each held-out dataset, the full preprocessing pipeline (HVG selection, PCA, scaling, Harmony fitting and random forest) was re-fitted on the training datasets only, to avoid any leakage from the held-out set. Held-out cells were projected onto the training-derived components and classified. For the glia-only transfer, the same procedure was applied to the oligodendrocyte + microglia subset. The number of evaluated stages equals the number of stages present in the held-out dataset: three for GSE277487 (control, acute and subacute); two for GSE160763 (control and subacute); and two for CEREBRI (control and acute, with its chronic-stage cells excluded from the balanced-accuracy metric and reported separately, as CEREBRI is the only dataset contributing the chronic stage). Because apparent subacute transfer can arise from classifiers collapsing onto the majority class, we additionally evaluated (i) subacute precision against the precision expected from a classifier assigning every cell to the subacute class (the subacute base rate in the held-out set), and (ii) one-vs-rest AUROC from class probabilities. These diagnostics were computed both on the three-dataset leave-one-out runs (using the stored confusion matrices) and in a stricter two-dataset replication (GSE277487 vs GSE160763 only, excluding the second CCI dataset from training, so that the held-out direction constitutes a pure cross-model test in which acute 6-h cells of GSE277487 were entirely unseen negative examples). In the two-dataset replication the classifier family was also varied (random forest, logistic regression, multilayer perceptron with two 64-unit hidden layers and early stopping, and 15-nearest neighbours) on identical train-only features, to test whether the collapse was model-dependent. To test whether the collapse was representation-dependent, the same transfer was repeated with a deep generative latent representation: a scVI model (30 latent dimensions) was fitted on the training glia only (raw counts of the train-only HVG genes) and held-out cells were mapped into the trained latent space without further training, before the identical random-forest classifier. Note that the acute training stage in the three-dataset runs derived from CEREBRI (24 h) whereas the held-out acute stage of GSE277487 is 6 h; immediate-early genes peak at ∼1–6 h, so the untransferability of the acute response reflects in part this time-point scheme mismatch. In addition to library-level evaluation, animal-level cross-validation was performed for the independent validation cohort (GSE247339): the cortex and hippocampus libraries of each animal were grouped together (GroupKFold over 12 animals), with significance assessed by 100 permutations of the animal–stage mapping (Supplementary Table S4). To verify the transcriptional state of the CEREBRI 24-h libraries used as the acute training stage, the library-level mean log1p-normalised expression of 15 canonical immediate-early genes (Fos, Fosb, Jun, Junb, Egr1, Egr2, Egr3, Arc, Nr4a1, Nr4a2, Nr4a3, Ier2, Dusp1, Npas4, Homer1) was computed across all cells of each CEREBRI library, comparing CCI-ipsilateral 24-h libraries (n = 6) with naïve control libraries (n = 10; Supplementary Fig. S2a).

### SHAP attribution

To obtain gene-level importance, a separate random forest (200 trees, maximum depth 15, minimum samples per leaf 10) was trained on the scaled HVG expression matrix of GSE277487, with the four sampling time points (control, 6 h, 2 d and 4 d) as classes. Mean absolute SHAP values were computed with TreeSHAP on a random subsample of 1,500 cells (deterministic seed).

### Cross-dataset gene-programme analysis

To ask which molecular components of the subacute response are reproducible across models, we computed, within microglia of each dataset, the library-level mean log1p-normalised expression difference between subacute-stage libraries (2–4 d in GSE277487; 7 d in GSE160763 and GSE247339) and control libraries, for every gene shared between datasets. A curated set of canonical programmes (complement/innate immunity; lysosome/phagocytosis; MHC-I/antigen presentation; iron/oxidative stress; lipid/cholesterol; a damage-associated-microglia (DAM)-like core of 13 genes (Trem2, Apoe, Ctsb, Ctsd, Ctsl, Ctsz, Lyz2, Spp1, Cd9, Gpnmb, Itgax, Lpl, Timp2) selected a priori from the DAM literature; and immediate-early genes as a negative control) was then tested for concordant direction of change across the CCI (GSE277487) and FPI (GSE160763) datasets. Significance was assessed against expression-matched permutation nulls: for each programme gene, 1,000 random genes were drawn from the same control-expression bin (20 quantile bins over the mean control expression across the two datasets), preserving the expression-level dependence of the effect sizes. Programme-level statistics reported are the concordance rate and the mean two-dataset delta, each against its matched null. The mild-FPI cohort (GSE247339) was examined as a third, independent test (acute 24-h, subacute 7-d and chronic 21-d libraries each versus the 21-d sham control libraries). All comparisons are library-level descriptive statistics; the small number of libraries (3 control and 6 subacute in GSE277487; 1 and 2 in GSE160763) precludes library-level hypothesis tests, and no cell-level statistics are reported for these analyses.

## Data availability

Raw data: GEO GSE277487, GSE160763, GSE247339 and GSE269748 (CEREBRI; CCI subset: control, 24 h, 6 months; 39,026 nuclei). Analysis scripts and result tables are available from the corresponding author upon reasonable request.

## Figure Legends

**Supplementary Fig. S1.**
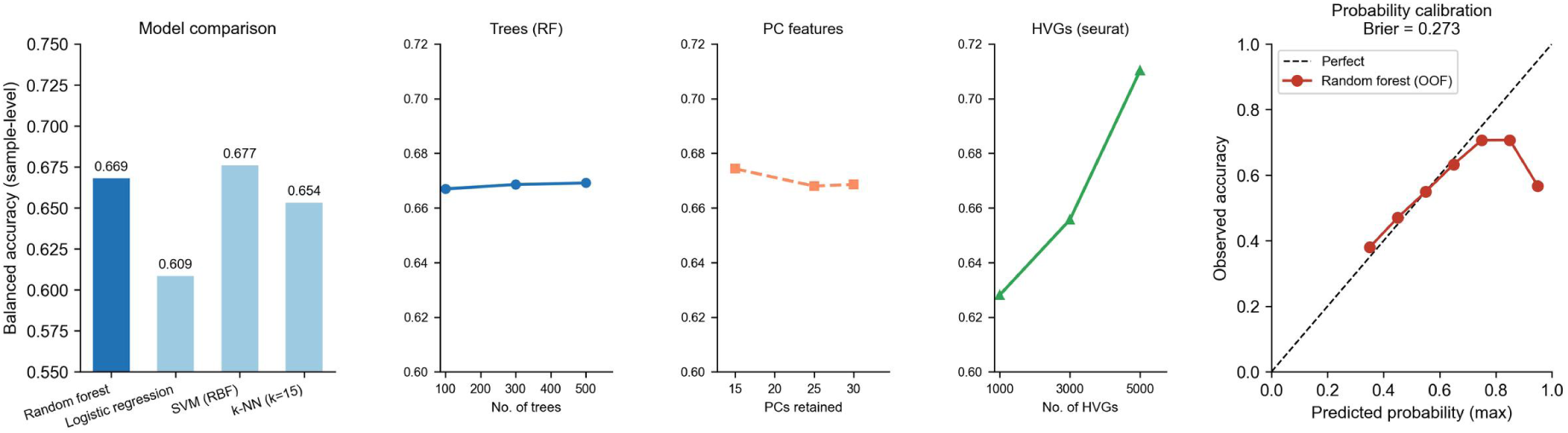
Robustness of the within-dataset classifier to model choice, hyperparameters and probability calibration (GSE277487). (a) Sample-level balanced accuracy of the four classifiers evaluated under the identical library-stratified cross-validation scheme: support vector machine with RBF kernel 0.677, random forest 0.669, k-nearest neighbours (k = 15) 0.654 and logistic regression 0.609. (b) Random-forest balanced accuracy as a function of hyperparameters, shown in three sub-panels with independent axes: number of trees (100–500; flat response), number of principal components retained (15– 30; flat response) and number of highly variable genes (1,000–5,000, seurat selection flavour; 0.63–0.71, modest upward trend). (c) Reliability diagram for out-of-fold maximum-class probabilities of the random forest (Brier score 0.273, versus 0.67 for a three-class chance baseline); grey dashed line, perfect calibration.

**Supplementary Fig. S2.** Stage-resolved expression in GSE277487 of the top-ranked SHAP genes of Fig. 5 that were detectable in this dataset (n = 47 of 50; 12 libraries). Rows are ordered by mean |SHAP| (top to bottom) and show row-wise z-scored mean log1p-normalised expression across control, 6 h, 2 d and 4 d samples. Immediate-early genes (Egr1, Fos, Fosb, Junb) peak at 6 h, whereas complement/innate-immunity, lipid/cholesterol, iron and astrocytic genes rise at the subacute (2–4 d) stages, specifying the pathological stage to which each gene’s attribution points.

## References

1. Maas AIR, Menon DK, Manley GT, et al. Traumatic brain injury: progress and challenges in prevention, clinical care, and research. Lancet Neurol. 2022;21(11):1004–1060. PMID: 36183712.

2. Jha RM, Rajasundaram D, Sneiderman C, et al. A single-cell atlas deconstructs heterogeneity across multiple models in murine traumatic brain injury and identifies novel cell-specific targets. Neuron. 2024;112(18):3069–3088.e4. PMID: 39019041.

3. Yang Q, Zhang L, Li M, et al. Single-nucleus transcriptomic mapping uncovers targets for traumatic brain injury. Genome Res. 2023;33(10):1818–1832. PMID: 37730437.

4. Izzy S, Liu Q, Fang Z, et al. Time-dependent changes in microglia transcriptional networks following traumatic brain injury. Front Cell Neurosci. 2019;13:307. PMID: 31440141.

5. Du QX, Li N, Dang LH, et al. Temporal expression of wound healing-related genes inform wound age estimation in rats after a skeletal muscle contusion: a multivariate statistical model analysis. Int J Legal Med. 2020;134(1):273–282. PMID: 30631906.

6. Lee H, Lee EJ, Park K, et al. MicroRNA transcriptome analysis for post-mortem interval estimation. Forensic Sci Int. 2025;370:112473. PMID: 40250071.

7. Hostiuc S, Rusu MC. The dynamics of neuroinflammation in traumatic brain injury: molecular markers useful for establishing the post-traumatic interval in forensic practice. Int J Mol Sci. 2026;27(4):2049. PMID: 41752185.

8. Tang X, Sumra V, Sato C, et al. The study of epigenetic clocks in former professional contact sports athletes with repetitive head impacts. J Neurol Neurosurg Psychiatry. 2026;97(9):827–833. PMID: 42114983.

9. Kunikeyev A, Salybekov AA, Yerimbetova A, Omarov B. Integrating mutation-derived and expression features from single-cell RNA sequencing: pitfalls of standard cross-validation in small-cohort settings. Int J Mol Sci. 2026;27(14):6429. PMID: 42511772.

10. Witcher KG, Bray CE, Chunchai T, et al. Traumatic brain injury causes chronic cortical inflammation and neuronal dysfunction mediated by microglia. J Neurosci. 2021;41(7):1597–1616. PMID: 33452227.

11. Korsunsky I, Millard N, Fan J, et al. Fast, sensitive and accurate integration of single-cell data with Harmony. Nat Methods. 2019;16(12):1289–1296. PMID: 31740819.

12. Ayerra L, Shumilov K, Ni A, et al. Chronic traumatic brain injury induces neurodegeneration, neuroinflammation, and cognitive deficits in a T cell-dependent manner. Brain Res. 2025;1850:149446. PMID: 39800094.

13. Zhang G, Diamante G, Ahn IS, et al. Thyroid hormone T4 mitigates traumatic brain injury in mice by dynamically remodeling cell type specific genes, pathways, and networks in hippocampus and frontal cortex. Biochim Biophys Acta Mol Basis Dis. 2024;1870(8):167344. PMID: 39004380.

14. Wang L, Ouyang D, Li L, et al. TREM2 affects DAM-like cell transformation in the acute phase of TBI in mice by regulating microglial glycolysis. J Neuroinflammation. 2025;22(1):6. PMID: 39800730.

15. Xia J, Sun CY, Su T, Xia SY, Zhao SZ, Zhang YM. Triggering receptor expressed on myeloid cell 2-mediated microglial phagocytosis in the subacute phase of traumatic brain injury exacerbates impaired memory. J Neurotrauma. 2026;43(3-4):317–340. PMID: 41700019.

16. Gao H, Di J, Clausen BH, et al. Distinct myeloid population phenotypes dependent on TREM2 expression levels shape the pathology of traumatic versus demyelinating CNS disorders. Cell Rep. 2023;42(6):112629. PMID: 37289590.

17. Li X, Xu Z, Fang J, Huang HD. P75NTR blockading inhibits Trem2+ M1 phenotype microglia activation and myelin damage following mild traumatic brain injury. Front Neurosci. 2025;19:1641112. PMID: 41574049.

18. Ciechanowska A, Ciapała K, Pawlik K, et al. Initiators of classical and lectin complement pathways are differently engaged after traumatic brain injury — time-dependent changes in the cortex, striatum, thalamus and hippocampus in a mouse model. Int J Mol Sci. 2020;22(1):41. PMID: 33375205.

19. He Z, Dony L, Fleck JS, et al. An integrated transcriptomic cell atlas of human neural organoids. Nature. 2024;635(8039):690–698. PMID: 39567792.

