## Supplementary Table S1 for "Conserved microglial programmes, non-transferable classifiers: boundaries of the single-cell TBI molecular clock"

Leave-one-dataset-out (LODO) cross-dataset transfer of post-injury stage classification: per-class recall, precision and one-vs-rest AUROC. All preprocessing (HVG selection, SVD, Harmony and the classifier) was fitted on training datasets only. Balanced accuracy (bacc) is computed over classes present in the held-out dataset (known classes); cells of a stage with no counterpart in the training data are reported separately as "unseen". Base-rate precision = precision expected from a classifier assigning every held-out cell to the subacute class. OVR AUROC = one-vs-rest AUROC for the subacute class (chance 0.5).

### Scheme 1: all cells, strict train-only preprocessing

| Held-out dataset | Evaluated stages | n_known_test | bacc | control recall | acute recall | subacute recall | chronic (unseen) |
| --- | --- | --- | --- | --- | --- | --- | --- |
| GSE277487 (CCI) | 3 (control, acute, subacute) | 162,892 | 0.310 | 0.328 | 0.002 | 0.599 | — |
| GSE160763 (FPI) | 2 (control, subacute) | 20,642 | 0.198 | 0.222 | — | 0.174 | — |
| CEREBRI (CCI snRNA) | 2 (control, acute) | 22,986 | 0.207 | 0.388 | 0.026 | — | n=16,040, 80.3% → subacute |

### Scheme 2: glia only (oligodendrocytes + microglia), strict train-only preprocessing — collapse diagnostics

| Held-out | stages | n_known_test | bacc | subacute recall | subacute precision | base-rate precision | % predicted subacute | control recall | acute recall |
| --- | --- | --- | --- | --- | --- | --- | --- | --- | --- |
| GSE277487 (CCI) | 3 | 30,046 | 0.338 | 0.913 | 0.513 | 0.501 | 89.2% | 0.093 | 0.008 |
| GSE160763 (FPI) | 2 | 12,940 | 0.473 | 0.748 | 0.584 | 0.583 | 74.6% | 0.198 | — |
| CEREBRI (CCI snRNA) | 2 | 17,524 | 0.037 | — | — | — | 88.2% | 0.045 | 0.029 |

*Subacute precision is indistinguishable from the subacute-only (base-rate) classifier in both cross-model directions, and 75–89% of all held-out glia are predicted subacute: the high recall reflects classifier collapse onto the majority class, not transferable discrimination.*

### Scheme 3: glial transfer from the CCI cohort to the independent mild-FPI cohort (GSE247339)

| Test | n_known_test | bacc | subacute recall | subacute OVR AUROC |
| --- | --- | --- | --- | --- |
| CCI-trained glial classifier on GSE247339 (7-d subacute vs control + 24-h) | 26,275 | 0.308 | 0.602 | 0.367 |

*21-d (chronic) cells of GSE247339 have no counterpart in the CCI training data and were excluded from the metrics.*

### Scheme 4: two-dataset strict cross-model replication (GSE277487 ↔ GSE160763; no CEREBRI in training) — classifier- and representation-independence

*Features identical to Scheme 2 (train-only HVG3000 + SVD30) unless stated; only the classifier/representation varies. In the FPI-trained → CCI-tested direction, acute (6 h) cells of GSE277487 were entirely unseen negative examples (FPI training data contain control and 7-d only). scVI: deep generative latent representation (30 dimensions) fitted on training glia only; held-out cells mapped into the trained latent space without further training.*

| Direction | Classifier / representation | bacc | subacute recall | subacute OVR AUROC | % predicted subacute |
| --- | --- | --- | --- | --- | --- |
| FPI → CCI | Random forest (SVD30) | 0.470 | 0.887 | 0.380 | 92.8% |
|  | Logistic regression (SVD30) | 0.489 | 0.907 | 0.466 | 93.3% |
|  | MLP 64×64 (SVD30) | 0.443 | 0.721 | 0.378 | 81.5% |
|  | k-NN k=15 (SVD30) | 0.463 | 0.872 | 0.356 | 92.0% |
|  | Random forest on scVI latent30 | 0.496 | 0.965 | 0.447 | 97.2% |
| CCI → FPI | Random forest (SVD30) | 0.460 | 0.899 | 0.544 | 90.2% |
|  | Logistic regression (SVD30) | 0.458 | 0.917 | 0.544 | 92.5% |
|  | MLP 64×64 (SVD30) | 0.337 | 0.586 | 0.475 | 60.0% |
|  | k-NN k=15 (SVD30) | 0.471 | 0.934 | 0.499 | 94.4% |
|  | Random forest on scVI latent30 | 0.500 | 0.983 | 0.536 | 98.3% |

### Notes

1. Chance levels: 1/3 for the three-stage GSE277487 direction; 1/2 for the two-stage directions (GSE160763, CEREBRI); 0.5 for OVR AUROC throughout.
2. The acute immediate-early response was essentially untransferable (acute recall 0.002 all cells; 0.008 glia, GSE277487 direction). The acute training stage derived from CEREBRI (24 h) whereas the held-out acute stage of GSE277487 is 6 h; immediate-early genes peak at ~1–6 h, so part of this failure reflects a time-point scheme mismatch.
3. High subacute recall in glia-only transfer (0.913 / 0.748) does not indicate a conserved transferable subacute state: precision equals the subacute-only baseline and OVR AUROC is at or below chance (Schemes 2 and 4), for four classifier families and for a deep generative (scVI) representation. The conserved cross-model component of the subacute response is a microglial gene-expression programme (DAM-like core; Supplementary Table S3), not a classifier state.
4. CEREBRI is the only dataset contributing the chronic (6-month) stage; its chronic cells are excluded from bacc. They were predominantly assigned to the subacute class (80.3% all cells; 91.2% glia). Because the LODO classifiers generally prefer the subacute class (Schemes 2 and 4), part of this assignment is a classifier-collapse artefact; the residual persistent-subacute-like interpretation is a hypothesis (Discussion).
5. CEREBRI (snRNA-seq) transfer failed overall (bacc 0.207 all cells; 0.037 glia), indicating the strongest platform-related shift of the three held-out datasets.
6. The loose scheme (Harmony fitted jointly before splitting) performed inconsistently (0.059–0.311 vs 0.198–0.310 strict), consistent with joint integration removing dataset-specific but time-informative variation; strict train-only estimates are primary.
7. A glial classifier trained on CCI recognised subacute (7-d) cells of the mild-FPI cohort with recall 0.602, but OVR AUROC was 0.367 (Scheme 3) — again collapse-dominated, not transferable discrimination.
