## Supplementary Table S2 for "Conserved microglial programmes, non-transferable classifiers: boundaries of the single-cell TBI molecular clock"

Marker gene sets used for cell-type annotation by data-driven marker scoring.

**Scoring rule.** For each cell, the score of a cell type was computed as the mean log1p-normalised expression (normalise_total to 10^4, then log1p) of the type's positive marker genes. A cell was assigned the type with the highest score, provided that score was positive; cells in which every marker score was zero were labelled Unknown. Marker sets are canonical genes for mouse brain cell types and were identical across all datasets and analyses.

| **Cell type** | **Marker genes** |
| --- | --- |
| Excitatory neuron | Slc17a7, Rbfox3, Neurod6, Tbr1, Camk2a |
| Inhibitory neuron | Gad1, Gad2, Sst, Pvalb, Npy |
| Astrocyte | Gfap, Aqp4, Slc1a3, Aldh1l1 |
| Microglia | C1qb, Tyrobp, Cx3cr1, Hexb, Aif1 |
| Oligodendrocyte | Mbp, Plp1, Mog, Mag |
| Oligodendrocyte precursor cell (OPC) | Pdgfra, Olig1, Cspg4 |
| Endothelial cell | Pecam1, Cldn5, Flt1 |
| Pericyte | Pdgfrb, Rgs5, Vtn |
| Unknown | fallback label: all marker scores zero |
