## Supplementary Figure S2 & Table S4 for "Conserved microglial programmes, non-transferable classifiers: boundaries of the single-cell TBI molecular clock"

### Supplementary Figure S2. Immediate-early gene expression in CEREBRI libraries

CEREBRI (GSE269748) CCI-ipsilateral 24-h libraries (n = 6, male) show no activation of canonical immediate-early genes relative to naïve control libraries (n = 10). Library-level mean log1p-normalised expression of 15 immediate-early genes (Fos, Fosb, Jun, Junb, Egr1, Egr2, Egr3, Arc, Nr4a1, Nr4a2, Nr4a3, Ier2, Dusp1, Npas4, Homer1) across all cells after QC (min 200 genes, <20% mitochondrial reads), log1p normalisation to 10,000 counts per cell. Control: 1.015 ± 0.197 (mean ± SD across libraries); CCI 24 h: 0.902 ± 0.072; difference −0.113. Points = individual libraries; error bars = SEM. The 24-h libraries used as the acute training stage therefore contained no immediate-early signal, supporting the time-point scheme mismatch interpretation of the untransferable acute response (Results; Methods).

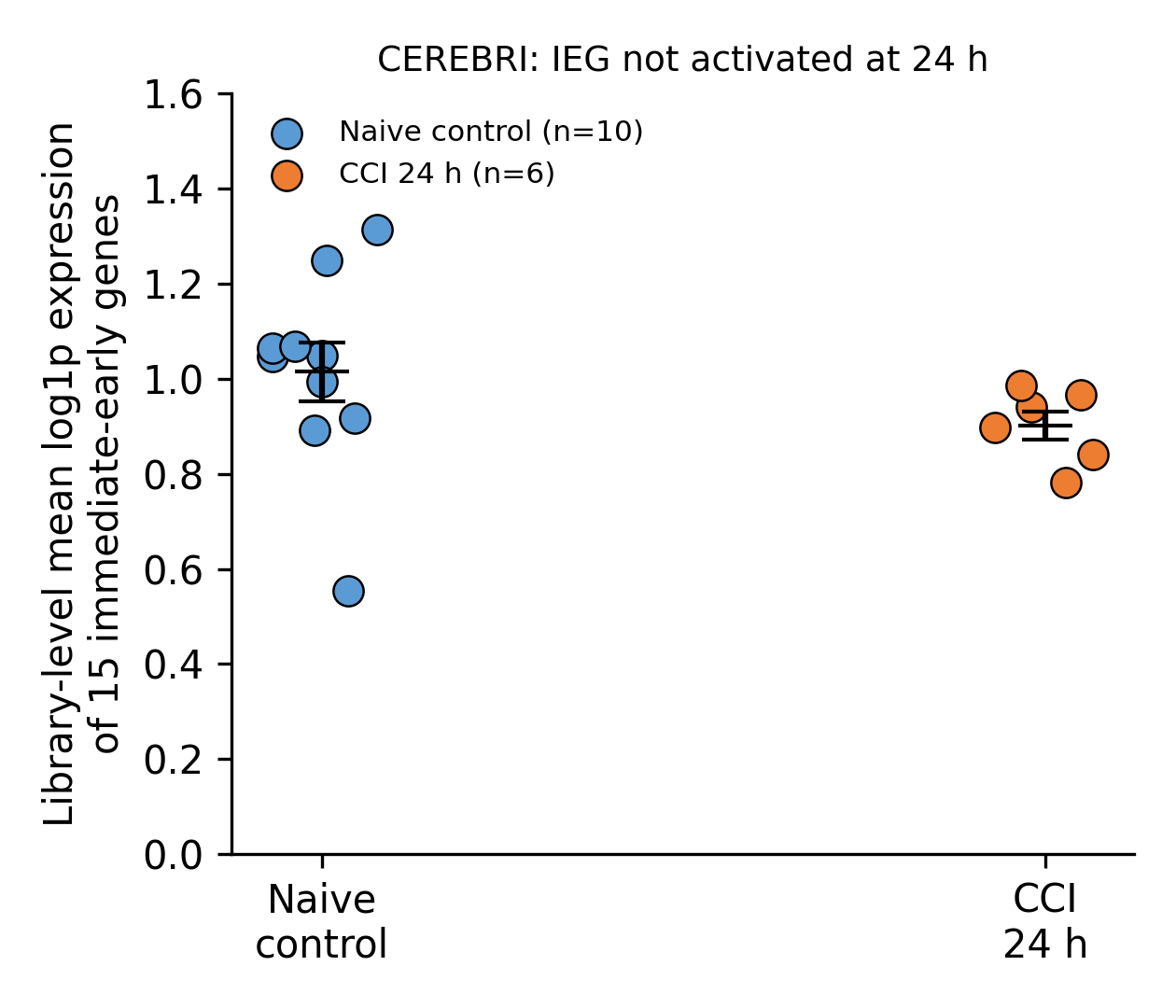

### Supplementary Table S4. Animal-level cross-validation of the independent validation cohort (GSE247339)

Cortex and hippocampus libraries of each animal were grouped together (GroupKFold over 12 animals; one library per region per animal, 24 libraries). The permutation null was generated by shuffling the animal–stage mapping 100 times (RF, 100 trees per permutation), preserving the number of animals per stage. For comparison the library-level (24-group) estimate is shown. Staging signal remains significant after removing within-animal correlation between regions.

| Evaluation level | Balanced accuracy | Permutation null (n = 100) |
| --- | --- | --- |
| Library-level (24 libraries) | 0.40 | — (1,000-permutation null: 0.19 ± 0.05; p = 0.0010; Methods) |
| Animal-level (12 animals) | 0.351 | 0.126 ± 0.049 (max 0.265); p = 0.0099; Cohen's d = 4.57 |
| Per-fold animal-level | 0.409 / 0.520 / 0.355 / 0.266 / 0.206 | — |
| Chance (4 stages) | 0.25 | — |
