## Supplementary Table S3 for "Conserved microglial programmes, non-transferable classifiers: boundaries of the single-cell TBI molecular clock"

Cross-dataset gene-programme analysis in microglia. Library-level mean log1p-normalised expression difference, subacute minus control (2–4 d in GSE277487; 7 d in GSE160763 and GSE247339). CCI = GSE277487 (snRNA-seq); FPI = GSE160763 (scRNA-seq); mTBI = GSE247339 (mild FPI; sham harvested at 21 d only). Significance against expression-matched permutation nulls (1,000 draws per gene from the same control-expression bin). Descriptive library-level statistics (library numbers are small; see Methods).

| Programme (a priori gene set) | n genes | CCI Δ | FPI Δ | mTBI Δ | Concordant CCI/FPI | matched-null p (concordance) | matched-null p (mean Δ) |
| --- | --- | --- | --- | --- | --- | --- | --- |
| complement/innate immunity | 8 | +0.652 | +0.040 | −0.344 | 7/8 | 0.082 | 0.001 |
| lysosome/phagocytosis | 14 | +0.560 | +0.104 | −0.161 | 13/14 | 0.008 | 0.001 |
| MHC-I/antigen presentation | 8 | +0.138 | +0.135 | −0.239 | 6/8 | 0.204 | 0.015 |
| iron/oxidative stress | 9 | +0.189 | +0.161 | −0.065 | 6/9 | 0.471 | 0.006 |
| lipid/cholesterol | 8 | +0.389 | +0.205 | −0.053 | 6/8 | 0.245 | 0.001 |
| DAM-like core (13 genes) | 13 | +0.507 | +0.130 | −0.130 | 13/13 | 0.004 | 0.001 |
| immediate-early (negative control) | 7 | −0.010 | +0.139 | −0.675 | 4/7 | n.d. | n.d. |

### DAM-like core, per-gene library-level deltas (13-gene a priori set)

| Gene | CCI Δ | FPI Δ | Concordant |
| --- | --- | --- | --- |
| Trem2 | +0.948 | +0.000 | yes |
| Apoe | +1.109 | +0.424 | yes |
| Ctsb | +0.633 | +0.264 | yes |
| Ctsd | +1.147 | +0.201 | yes |
| Ctsl | +0.718 | +0.110 | yes |
| Ctsz | +0.913 | +0.071 | yes |
| Lyz2 | +0.284 | +0.153 | yes |
| Spp1 | +0.145 | +0.070 | yes |
| Cd9 | +0.287 | +0.173 | yes |
| Gpnmb | +0.035 | +0.002 | yes |
| Itgax | +0.014 | +0.019 | yes |
| Lpl | +0.057 | +0.011 | yes |
| Timp2 | +0.304 | +0.197 | yes |

### Library-level DAM13 score (mean of 13 genes), control vs subacute

| Dataset | control libraries (mean ± SD) | subacute libraries (mean ± SD) | Cohen's d |
| --- | --- | --- | --- |
| CCI (GSE277487) | 3 (0.50 ± 0.11) | 6 (1.02 ± 0.17) | 3.2 |
| FPI (GSE160763) | 1 (1.57) | 2 (1.70 ± 0.00) | n.a. (single control library) |
| mTBI (GSE247339) | 6 (1.07 ± 0.08) | 6 (0.94 ± 0.15) | −1.1 |

1. The FPI cohort shows the same direction but much smaller amplitude, and its microglial signal is carried largely by myelin-phagocytic (Mbp+) cells; the single control library precludes effect-size estimation.
2. In the mild-FPI cohort the programme was not up-regulated at any sampled time point (24 h, 7 d, 21 d; 21-d TBI vs 21-d sham p = 0.31). A TREM2-positive microglial phenotype has nonetheless been described after mild TBI (main text ref. 17), so this negative result is cohort-design-specific.
3. Immediate-early genes serve as a negative control (acute-peaking module, not expected in the subacute window).
